# On the typical development of the central sulcus in infancy: a longitudinal evaluation of its morphology and link to behaviour

**DOI:** 10.1101/2025.02.19.639011

**Authors:** Amaia Dornier, Alexia Gérard, Yann Leprince, Lucie Hertz-Pannier, Jean-François Mangin, Marianne Barbu-Roth, Jessica Dubois, Dollyane Muret

## Abstract

**Introduction:** The progressive folding of the cortex is an important feature of neurodevelopment starting early during gestation. The central sulcus (CS) is one of the first to fold. Since it represents the anatomical boundary between primary somatosensory and motor functional regions, its developing morphology may inform on the acquisition of sensorimotor skills. We aimed to identify potential asynchronous morphological changes along the CS during infancy, with the hypothesis that this may reflect the gradual emergence of body usage.

**Method:** Based on 3T anatomical magnetic resonance imaging (MRI) and dedicated post-processing, we characterized the evolution in CS depth and curvature in 33 typical infants (aged 1 and 3 months, 22 with longitudinal data) in relation to 23 young adults as a reference. Four regions of interest (ROIs) along the CS, supposed to correspond to different parts of the body and one centred on the hand knob (HK), were reproducibly examined and compared across groups. We also explored the relationship between the age-related changes in morphological features and the global motor scaled scores evaluated at 3 months of age with the Bayley Scales of Infant and Toddler Development.

**Results:** While all ROIs showed significant increases in CS depth and curvature between 3-month-olds and adults, the results were more variable between 1 and 3 months of age depending on cross-sectional and longitudinal analyses. The central-medial and central-lateral regions showed the most consistent increase in depth. Besides, motor development at 3 months of age was not significantly related to CS morphological changes, but a positive trend was observed for depth changes in the (HK-related) central-medial ROI.

**Conclusion:** The rapid evolution of CS folding during infancy may reflect the intense but asynchronous maturation of the brain sensorimotor system, with the differential growth of cortical areas related to body parts and underlying white matter connections. Although it will have to be replicated on larger groups and at other ages, this longitudinal and multimodal study highlights the potential of characterizing CS features as key markers of early sensorimotor development, both at the cerebral and behavioural levels. Combining anatomical and functional neuroimaging could provide deeper insights into the relationship between CS morphology and somatotopic organization in typical infants, but also in infants at risk of developing motor disorders.

## Introduction

The development of the human brain is an extraordinarily intricate and complex process that relies on a series of key cellular mechanisms beginning during gestation (see [1] for a review). At the macroscopic level, an important feature of brain development lies in its progressive folding. Brain folding begins around the 14^th^ week of gestation with the formation of primary sulci, followed by secondary folds appearing around the 32^nd^ week, and tertiary folds beginning to form from the 38^th^ week [1, 2], resulting in the well-defined sulci and gyri typical of the mature brain.

Among early developing sulci, the central sulcus (CS) is one of the first to form since its first dimples are visible from the 18^th^ week, with an earlier onset in the right hemisphere than in the left [1, 3]. The CS is a critical marker of brain organization as it delineates the frontal and parietal lobes and marks the boundary between the primary motor (M1) and somatosensory (S1) cortices hosting sensorimotor functions. Interestingly, its shape features in adults have recently been linked to its topographical organization [4] that characterizes the well-known *Homunculus* first described in humans by Penfield and Boldrey [5]. In particular, the “hand-knob” is a key marker of hand sensorimotor function: its prominent asymmetrical morphology and central position in the CS are thought to reflect mature motor abilities and lateralization to some extent, notably with regard to handedness and speech-related functions in adults [6]. In addition, the early shape features of the CS observed in preterm infants post-natally were suggested to help predicting their motor abilities and manual laterality at 5 years of age [7]. Together, this suggests that the CS could be used as an early anatomical marker of sensorimotor development, a function that has proved to be challenging to study at the functional level in young infants [8, 9]. However, the development of the CS and the evolution of its morphological features in the first months after birth remain poorly understood.

Considering the CS as a whole, some of its shape characteristics were found to be already encoded at 30 weeks of post-menstrual age (PMA) in premature infants imaged close to birth [7]. Its folding then becomes more elaborate in the following weeks, as observed in the same infants imaged at term-equivalent age, and some asymmetries in shape were shown between the left and right hemispheres [7]. Interestingly, a quantitative analysis of the CS depth in early childhood revealed different variations with age depending on the location on the CS: while the sulcus depth showed a rightward asymmetry at 1 year of age in the lateral part of the CS, this disappeared before 2 years of age [10]. Besides, while the CS demonstrated an overall increase in depth between 1 and 3 years of age, this was less marked in the central region of the sulcus [10]. The hypotheses put forward to explain these heterogeneous changes in depth along the CS involved respectively the development of language skills for the lateral part of the CS, and the formation of a’*Pli de Passage’* [11] in the central region, consequently to the development and myelination of underlying white matter connections [10]. However, while unexplored so far, the first post-natal months are an important period not only for brain folding [12] but also for the intense sensorimotor experiences and skill acquisitions emerging at these ages [9, 13, 14].

In addition to genetic factors, these behaviours and experiences are thought to impact but also to rely on the development of underlying brain networks [7, 15]. In particular, the development of both behaviours and underlying brain systems is supposed to proceed asynchronously along the body axis, from proximal to distal regions and from superior to inferior regions [16]. Such asynchrony in maturation is a well-known phenomenon across cerebral networks [17, 18] and has been previously highlighted using magnetic resonance imaging (MRI) in typical infants, both at the white matter and cortical levels (e.g., [19, 20]). As such, one could expect different developmental trajectories of maturation along the CS as sensorimotor acquisitions and body usage emerge, but this has never been explored during infancy.

Leveraging the proven advantages of MRI to map the folding process non-invasively and quantitatively in early development (see [2] for a review), we aimed to fill this gap by imaging typical infants at 1 and 3 months of age, in association with behavioural assessment at the latter age. We examined the evolution of key morphological markers of the CS (depth, curvature) with reference to a group of young adults as these morphological features also vary along the CS in the mature brain. As we assumed the development of folding to proceed asynchronously along the CS, we considered different regions supposed to be associated with different body parts, in particular the hand-knob that was identified at the individual level. We further related sensorimotor acquisitions with the evolution of CS morphological markers in order to contribute to our understanding of how early brain structure supports behavioural development in infants.

## Methods

### Participants and data collection

#### Infant group

Thirty-three typically developing infants (16 girls) were included in this study. They were born of single full-term pregnancies (i.e., between 37.43 and 41.86 weeks of gestational age) and had no neurological risk or background (i.e., incident-free pregnancy and birth). MRI acquisition was performed on 26 of these infants (13 girls) at approximately 1 month (1M) of age (post-menstrual age PMA at MRI: mean ± standard deviation: 45.20 ± 0.65 weeks, range: 44.14–46.43 weeks), and on 29 of these infants (14 girls) at 3 months (3M) of age (PMA at MRI: 53.21 ± 0.82 weeks, 51.43–54.71 weeks). The dataset included 22 infants (11 girls) with longitudinal data collected at the two ages.

MRI examinations were performed using a 3 Tesla magnetic field scanner (Siemens Prisma) and a 20-channel head coil for most infants (except for 4 infants at 1M for whom a 64-channel head coil was used). Acquisitions were carried out during the infant’s spontaneous sleep, after nursing, swaddling and putting earplugs and headphones to protect from the scanner noise. For anatomical explorations with a good contrast between the developing grey and white matter, T2-weighted (T2w) images were acquired with a 2D Turbo Spin Echo sequence with the following parameters: TR = 4200 ms, TE = 149 ms, flip angle = 140°, parallel imaging with GRAPPA acceleration factor = 2, spatial resolution = 0.8×0.8×2 mm, 60 slices (acquisition time = 1min55s). Images were acquired in axial plane for all infants, and an additional scan was acquired in sagittal plane for 20 infants (7 at 1M, 15 at 3M).

Alongside the MRI acquisition, a behavioural evaluation of the infants was carried out on the same day, but in this study, we focused on the data collected at the age of 3M. The gross and fine motor (GM / FM) skills were assessed with the Bayley Scales of Infant and Toddler Development 3^rd^ edition (BSID-III; [21]). At 3M, up to the first 26 GM items and 21 FM items are evaluated ([22]), stopping after 5 consecutive failed items (i.e., child developmental ceiling). GM and FM scaled scores, ranging from 1 to 19, were obtained, providing information on the infant’s motor development in relation to other infants of the same age (calibration on a North American population). The global motor scaled score was computed as the sum of GM and FM scaled scores.

#### Adult group

This study also considered data from 23 healthy young adults from the Human Connectome Project (HCP), born full term (12 women, range of ages: 22-35 years) who were selected from the Human Connectome Project (HCP) database. None showed brain abnormality or had a history of diabetes, hypertension, excessive alcohol consumption, or substance abuse, nor any sibling with severe neurodevelopmental, neuropsychiatric, or neurological disorders. MRI data acquisition was performed using a 3 Tesla magnetic resonance scanner (Siemens Connectome Skyra) equipped with a 32-channel head coil. For anatomical explorations, T1-weighted (T1w) images were collected using a 3D gradient-echo sequence (TR = 2400 ms, TE = 3.16 ms, TI = 1000 ms, parallel imaging with GRAPPA acceleration factor = 2, isotropic resolution of 1 mm) [23]. The high quality of MRI data was one of the criteria to select these subjects, as well as data availability and pre-processing in our institution.

### Processing of MRI data

#### Infant group

For each infant at each age, axial T2w images were first Fourier-resampled to an isotropic spatial resolution of 0.8 mm. Segmentation processing was performed using iBEAT2 (version 120, [24]). After bias correction, a first segmentation was carried out to generate the brain mask, which was manually corrected if needed (for 5 participants). Within this adjusted brain mask, brain tissues (grey and white matter, cortico-spinal fluid) were segmented (iBEAT2 version 205). The resulting segmentations were further processed using BrainVISA software [25] through the “Morphologist” processing pipeline [26] to identify cortical sulci. After correcting the segmentations in the central regions when needed, the inner and outer cortical surfaces were reconstructed for both hemispheres. Objects representing cortical sulci were identified between the brain hull and pial surfaces, and automatically labelled [27]. Global brain volume was estimated as the sum of hemispheric volumes contained within the closed outer cortical surfaces [12, 27].

#### Adult group

T1w images were fully processed using BrainVISA and Morphologist pipeline [26], to obtain brain tissue segmentation, cortical surface reconstruction and identification of cortical sulci as done for infants data.

### Analysis of central sulcus morphology

#### Parametrization of central sulcus

Based on the whole brain labelled set of cortical sulci, the central sulcus (CS) of each participant was visually checked to ensure a proper and consistent reconstruction and labelling over infants and adults, particularly for longitudinal data. The length of left and right CS was estimated for each subject. The BrainVISA “Cortical Surface” pipeline [29], along with the sulcus parameterization toolbox [29, 30, 31], were further used to extract and parameterize the CS through the definition of a curvilinear coordinate (from 0 at its upper medial edge to 100 at its lower lateral edge, see Figure 1).

**Figure 1.**
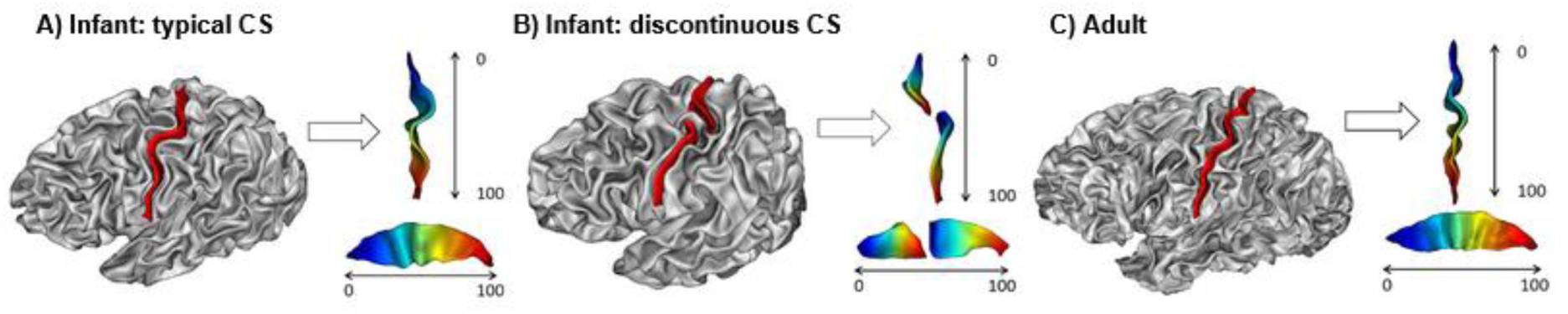
Examples of individual central sulcus (CS) and CS parameterization (left hemisphere) A) 1 month old infant showing a typical CS. B) 1 month old infant showing a discontinuous CS. C) Adult showing a typical CS. All CS are displayed (in red) on the cortical surface of the subject. CS are parameterized along the sulcus with coordinates ranging from 0 (most medial edge) to 100 (most lateral edge) and displayed from both a dorsal (top) and lateral (bottom) view to highlight the CS curvature and depth respectively. CS coordinates are coded by the gradient of colour: blue = 0, red = 100.

#### Measure of CS depth and curvature

Using BrainVISA toolboxes, each CS coordinate was associated with a depth value (i.e., geodesic distance between the crest and the bottom of the sulcus) and a sulcal profile value (i.e., signed distance from the CS plane of inertia that captures its main orientation). Zero sulcal profile values at coordinates 0 and 100 were replaced by neighbouring values at coordinates 1 and 99, and a one-dimensional Gaussian filter (sigma = 2) was applied to the sulcal profile values at all coordinates. The CS local curvature was then computed from the neighbouring sulcal profile values according to the following formula:

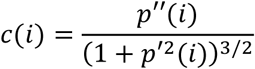

With 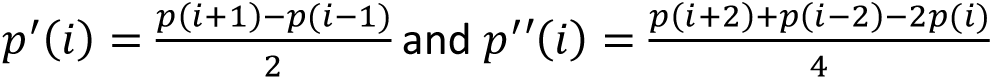

*c* the curvature, *p* the sulcal profile, and *i* the position along CS (2≤i≤98).

In order to inform on the local sulcal curviness regardless of its direction, we further considered the absolute curvature values.

#### Definition of regions of interest

For each participant (infant / adult), the coordinate of the centre of the hand-knob (HK) along the CS was identified based on anatomical (T2w / T1w) images and 3D surfaces, according to its shape and localization [32], checking for overall consistency across left and right hemispheres. Four regions of interest (ROIs) were then defined at the individual level: one was centred on the HK coordinate (hereafter, central-medial), two were placed respectively at the lowest (most medial) and highest (most lateral) coordinates of the CS (hereafter, medial and lateral), and a fourth ROI was placed at equidistance between the central-medial and lateral ROIs (hereafter, central-lateral). To minimize noise issues at CS extremities, the 5 lowest and highest coordinates of the CS were excluded before defining the ROIs. Based on these criteria and given the range of HK coordinates observed across both the infant and adult groups, a consistent ROI size of 14 coordinates was used for all subjects and for both hemispheres to obtain comparable ROIs across participants, though defined based on the individual’s anatomy.

Depth and absolute curvature values were measured for each hemisphere along the CS coordinates, as well as averaged over its entire length and over each ROI. Averages over left and right hemispheres were also considered.

### Statistical analyses

#### Outlier identification

Within each age group (1M, 3M, adults), subjects with an absolute z-score in brain volume higher than 2 were excluded from all analyses, as well as subjects with discontinuous (i.e., interrupted) CS. When analysing morphological features (depth, absolute curvature), subjects displaying z-scores higher than 2 for averaged measures (over left and right CS) were considered as outliers. As for analyses targeting behavioural scores, we further excluded subjects with FM and GM scores lower than or equal to 7 since this threshold has been associated with a risk of atypical neurodevelopment [21].

#### Evaluation of interhemispheric asymmetries

For each morphological feature *F* (i.e., depth and absolute curvature averaged over CS) and each subject, an asymmetry index (*AI)* was computed between the left (L) and right (R) hemispheres following this equation [33]:

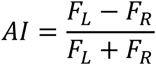

A positive index indicates left asymmetry, while a negative index indicates right asymmetry. The presence or absence of asymmetry was evaluated for each age group independently using Wilcoxon test on AI. Correction for multiple comparisons (three groups) was performed using False Discovery Rate (FDR) approach, and a statistical threshold p<0.05 was considered as significant. In case of non-significance, the average between right and left measures was considered for further analysis. Supplementary analyses on inter-hemispheric asymmetries (see Supplementary Information SI.1) were performed on the HK coordinates (see Supplementary Table 1.1) and on the morphological features obtained within each ROIs (see Supplementary Table 1.2).

#### Evolution and relationships between brain volume and CS measures

To assess whether the differences between groups were of the same order of magnitude between brain features, the means (noted 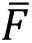) were compared between the 1M and 3M groups using the formula of relative difference: 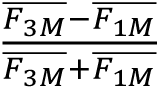(and similarly for the 3M and adult groups).

For each CS morphological feature (i.e., depth and absolute curvature averaged over the left and right CS), we further investigated the relationship with brain volume using Pearson correlations performed after logarithmic linearization to account for possible allometries [28]. This was done for each age group separately and when combining 1M and 3M infants. Correlations between averaged depth and absolute curvature were also tested for each group and combined infant groups.

#### Impact of age group and ROI localization on CS features

We then aimed to evaluate whether each CS morphological feature (i.e., depth, absolute curvature averaged over left and right hemispheres) differed across the four ROIs in a distinct way across the three age groups (1M, 3M, adults). First, a linear mixed-effects model (LMM) was performed on the entire dataset (both longitudinal and cross-sectional age groups, but excluding outliers identified for the considered feature within each group) considering Group and ROI as fixed-effect predictors, as well as their interaction (Group * ROI), and participants as random-effect predictors. Post-hoc tests were conducted using pairwise comparisons of estimated marginal means (EMMs), and adjustments for multiple comparisons were applied using the FDR approach.

Second, we focused on infants with longitudinal data to compute the relative change in each feature *F* of each ROI between 1M and 3M (in %) using the following equation:

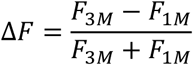

We tested whether this relative change was different from 0 using Student t-tests (four ROIs). The effect of ROIs was then assessed using an analysis of variance (ANOVA) with the factor ROI. Post-hoc tests consisted of paired t-tests across ROI pairs (6 comparisons). For each analysis, correction for multiple comparisons was performed with the FDR approach.

For both depth and absolute curvature, supplementary analyses along the CS were performed (see Supplementary Information SI.2 and Supplementary Figure 2.1).

#### Relationship between the evolution of CS morphology and sensorimotor development

To evaluate whether sensorimotor acquisitions at 3M were related to the developmental evolution of CS regions, we considered the global motor scaled score of infants and related it with longitudinal MRI measures (1M, 3M). Since the inter-individual variability in the age gap between 1M and 3M might impact the age-related difference in MRI features, we considered the variables 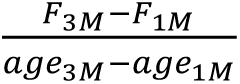 in linear models all describing the global motor scaled score but testing different hypotheses successively. The first model evaluated the relationship with a global brain feature not specific to CS, so the brain volume was considered. With the second model, we tested the relationship with global CS morphological characteristics, considering depth and absolute curvature averaged over the sulcus. The third model questioned a variable relationship depending on location along the CS: we considered the morphological characteristics averaged over the two central ROIs independently, as the medial and lateral ROIs showed less robust changes with age. For the last two models, we had checked beforehand that depth and absolute curvature were not correlated, and that they did not depend on brain volume changes. Correction for multiple comparisons was performed across models with FDR approach.

### Additional validation analyses

Supplementary analyses on the intra-subject variability related to the pipeline used for sulcus identification (see Supplementary Information SI.3, Supplementary Figure 3.1 and Supplementary Tables 3.1 and 3.2) and to the T2w reconstruction approach (super resolution vs. resampling; see Supplementary Information SI.3, Supplementary Figures 3.2 and 3.3 and Supplementary Tables 3.3 and 3.4) were performed.

## Results

### Global CS morphology

#### CS identification

Reliable CS reconstructions were obtained for all subjects (examples in Figure 1A and C) thanks to the careful verification of brain segmentations and manual corrections whenever needed. Among all subjects, three infants showed a discontinuous CS (example in Figure 1B): two at both ages (longitudinal data) in the left hemisphere, and one at 3 months in the right hemisphere. These infants were excluded from all analyses. As expected from the increasing brain size with age (Figure 2A), CS length also qualitatively increased from 1M to 3M and from 3M to adults (Figure 2B).

**Figure 2.**
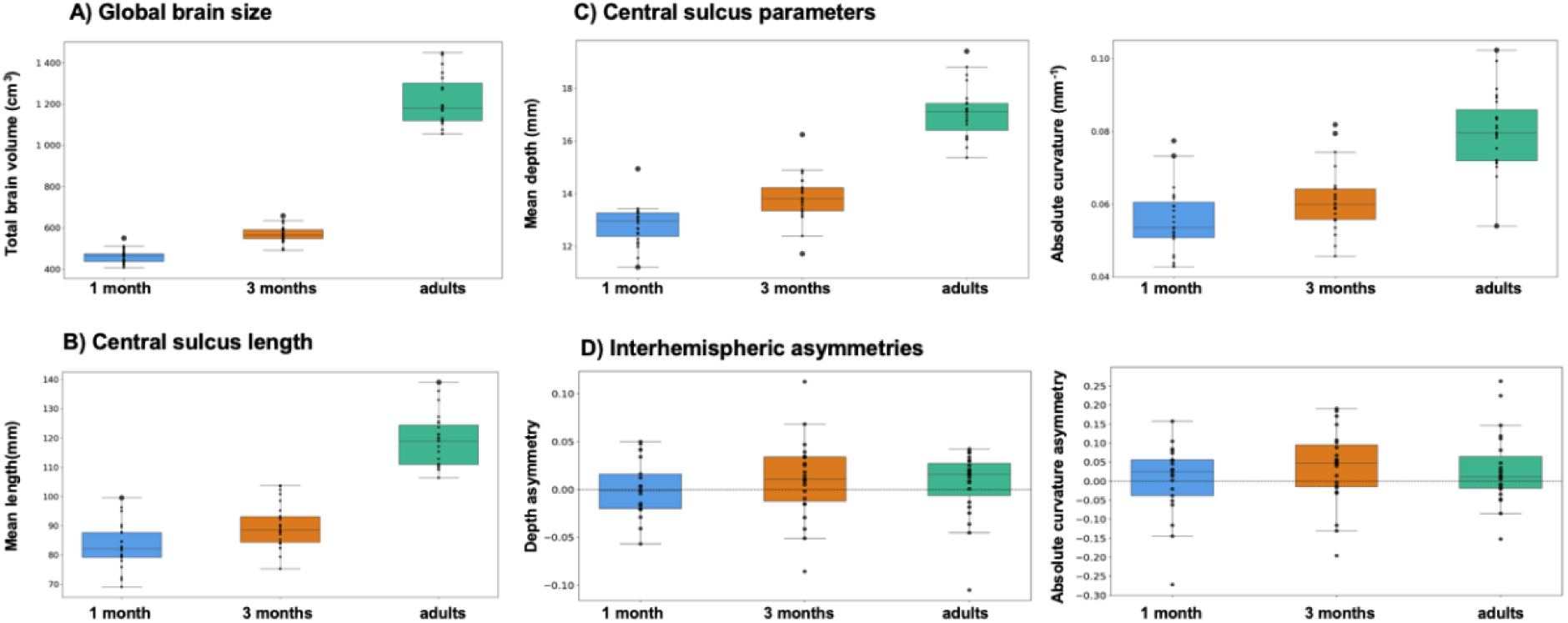
Quantification of brain features, outlier identification and evaluation of interhemispheric asymmetry within each group. A) Boxplot of the brain volume (in cm^3^) observed for each group (infants at 1 and 3 months of age and adults). Individual data points are represented on top of the boxplots. One infant was identified as an outlier at both ages (bigger points). B) Boxplot of the length (in mm) averaged over the left and right CS and observed for each group. C) Boxplots of the depth (left column) and absolute curvature (right column) averaged over the left and right CS and observed for each group (outliers shown with bigger points). D) Boxplots of the interhemispheric asymmetry index computed for each group for the average depth of the CS (left column) and the average absolute curvature of the CS (right column). A positive asymmetry indicates higher values in the left hemisphere relative to the right hemisphere (and vice versa). No significant asymmetries were observed.

#### Outlier identification

In terms of brain volume, we identified one outlier infant at both 1M and 3M, which was excluded from all analyses, and no adult outlier (Figure 2A). Regarding depth measures (averaged over the whole left and right CS), two outlier infants were identified at both ages, as well as one adult (Figure 2C left column). Regarding absolute curvature measures, one outlier infant at both 1M and 3M, one at 1M and one at 3M, along with two adults, were identified (Figure 2C right column). The outlier infants for depth were not the same as those for curvature, but the adult identified as outlier for depth was also an outlier for curvature. These subjects were excluded from the analyses of the corresponding feature, in addition to subjects with discontinuous CS or outlier brain volume, leading to the following numbers of subjects: depth: n = 21 (1M), 23 (3M), 22 (AD); curvature: n = 21 (1M), 23 (3M), 21 (AD).

#### Interhemispheric asymmetries

Regarding interhemispheric asymmetries, the Wilcoxon tests performed on asymmetry indices for depth and absolute curvature averaged over the whole CS showed no significant differences for any group of subjects (Figure 2D, Table 1). This suggested that overall CS morphological features were not different between the left and right hemispheres. Averaged left and right measures were thus considered in the following analyses.

**Table 1.** Summary statistics on interhemispheric asymmetries. Results of the Wilcoxon tests on asymmetry indices for depth (A) and absolute curvature (B) averaged over the CS, for the 1 month (1M), 3 months (3M), and adult (AD) groups.

| Group | N | median | W | p-value | BF10 |
| --- | --- | --- | --- | --- | --- |
| **A) Depth** |  |  |  |  |  |
| 1M | 21 | -0.001 | 110 | 0.865 | 0.23 |
| 3M | 23 | 0.01 | 95 | 0.20 | 0.42 |
| AD | 22 | 0.016 | 88 | 0.222 | 0.26 |
| **B) Absolute curvature** |  |  |  |  |  |
| 1M | 21 | 0.025 | 93 | 0.452 | 0.27 |
| 3M | 23 | 0.048 | 76 | 0.06 | 0.83 |
| AD | 21 | 0.012 | 77 | 0.191 | 0.79 |

#### Relationships between features

Based on the computation of the feature relative difference comparing mean values over groups, we observed larger increase in brain volume (1M-3M: 10.48%, 3M-AD: 36.53%) than in CS depth (1M-3M: 3.96%, 3M-AD: 10.20%) and absolute curvature (1M-3M: 5.16%, 3M-AD: 15.03%). Increases were also qualitatively larger between infants at 3M and adults than between both infant groups.

We then evaluated whether CS morphological features (depth and absolute curvature) were related to brain volume using Pearson correlations between measures on a logarithmic scale and over each group. No significant result was found for infants at 1M (depth: N=21, r=0.27; curvature: N=21, r=0.23; all p values > 0.30) or at 3M (depth: N=23, r=0.36; curvature: N=23, r=0.20; all p values > 0.30), but a significant correlation was observed when considering both groups together (depth: N=44, r = 0.70, p = 2.08.10^-7^; curvature: N=44, r = 0.45, p = 2.7.10^-3^). In adults, these correlations were also significant (depth: N=22, r = 0.69, p = 3.75.10^-4^; curvature: N=21, r = 0.62, p = 1.55.10^-3^). This suggests that greater brain volume may be associated with deeper and curvier CS across different stages of development.

Besides, no significant correlation was observed between CS depth and absolute curvature in each group (1M: N=19, r = 0.29; 3M: N=21, r = 0.34; AD: N=21, r = 0.33; all p values > 0.10), except when considering both infant groups together (N=40, r = 0.45, p = 0.003), suggesting a low association between CS depth and curvature.

### Regional CS morphological features

#### Hand knob identification

Hand knobs (HK) were visually identified in all subjects, whatever their age (Figure 3). The average position of the HK remained stable across ages, with minor variations between the left (coordinates: 41.8 ± 7.4, range: 27–59 (1M); 41.6 ± 7.7, range: 27–60 (3M); 43.7 ± 4.1, range: 30–49 (adults)) and right hemisphere (coordinates: 39.9 ± 5.5, range: 29–52 (1M); 38.9 ± 5.0, range: 30–47 (3M); 41.1 ± 4.8, range: 32–49 (adults); see Supplementary Information SI.1 and Supplementary Table 1.1 for additional analyses on inter-hemispheric asymmetries in HK location). On average over each group, HK were localized at the level of a local plateau in depth (the highest depth was observed at higher coordinates) and high absolute curvature (Figure 3).

**Figure 3.**
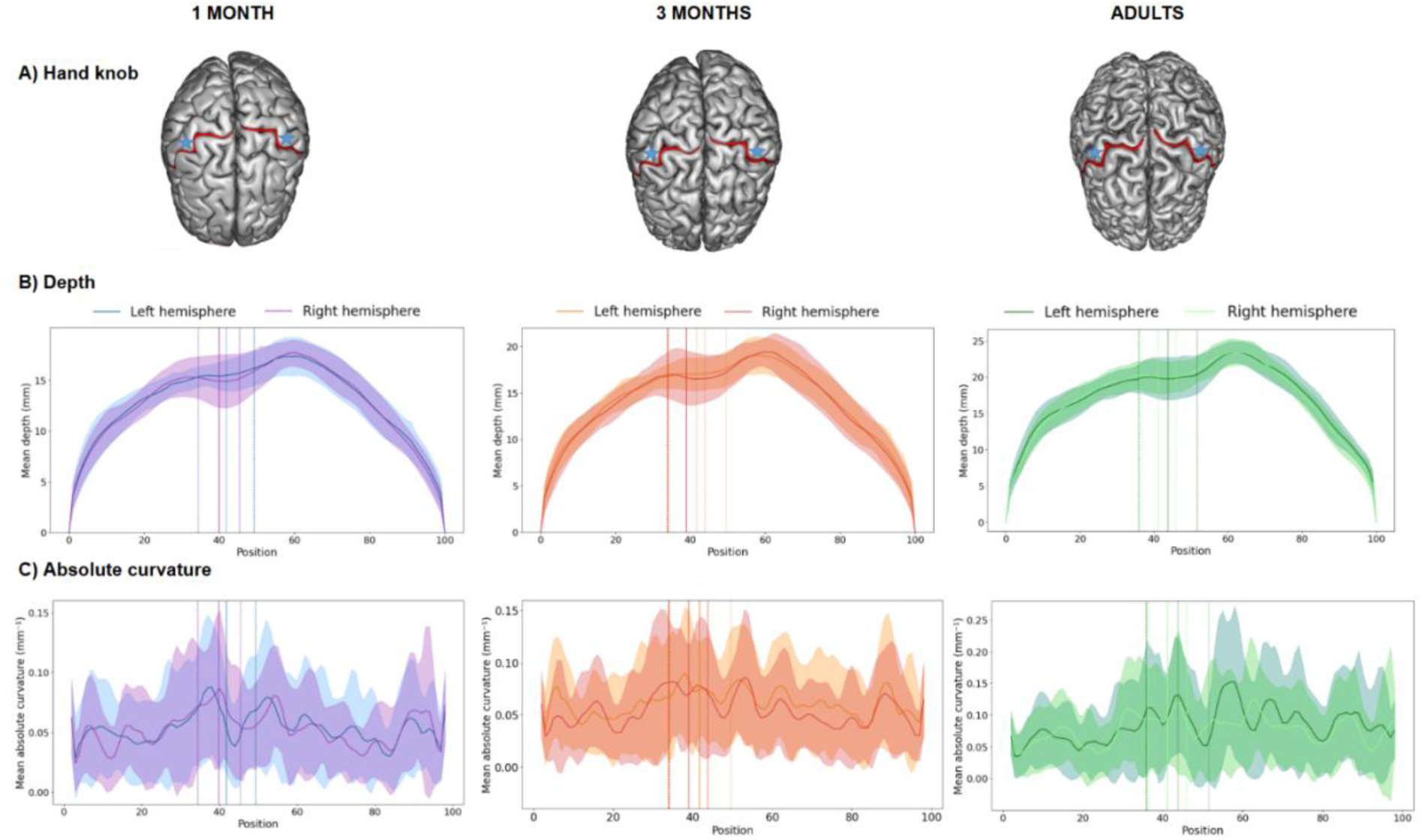
Identification of the hand knob along the central sulcus parametrization for each age group. A) Example of hand knob (HK) identification for an infant at 1 month (1M) and 3 months (3M) and an adult (AD). For each group, HK localizations (average shown with the vertical continuous line and standard deviation with dotted lines) are superimposed to CS parametrization (average shown with the continuous curves and standard deviations with the shaded area) for depth (B) and absolute curvature (C) for the left (dark colours) and right (light colours) hemispheres (average metrics over each group, with standard deviations in shaded colours).

#### Feature dependence on age group and ROIs localization

A qualitative comparison of CS parameterized curves first revealed some differences across age groups, with an increased depth and curvature from 1M to 3M infants and to adults in most coordinates of the sulcus but more particularly in the central part (Figure 4B; see Supplementary Information SI.2 and Supplementary Figure 2.1 for additional analyses along the CS). The HK identification allowed us to consider four ROIs along the CS (Figure 4A), with consistent averaged localization and specificities across groups (Figure 4B): the (HK-related) central-medial ROI showing pronounced curvature; the central-lateral ROI showing the highest depth; the medial and lateral ROIs at the CS edges showing lower depth and lower absolute curvature than central ROIs. In order to further explore the feature dependence on position along the CS and on age groups, CS morphological features were then averaged over each left and right ROIs defined at the individual level (see Supplementary Information SI.1 and Supplementary Table 1.2 for additional analyses on inter-hemispheric asymmetries in ROI depth and absolute curvature).

**Figure 4.**
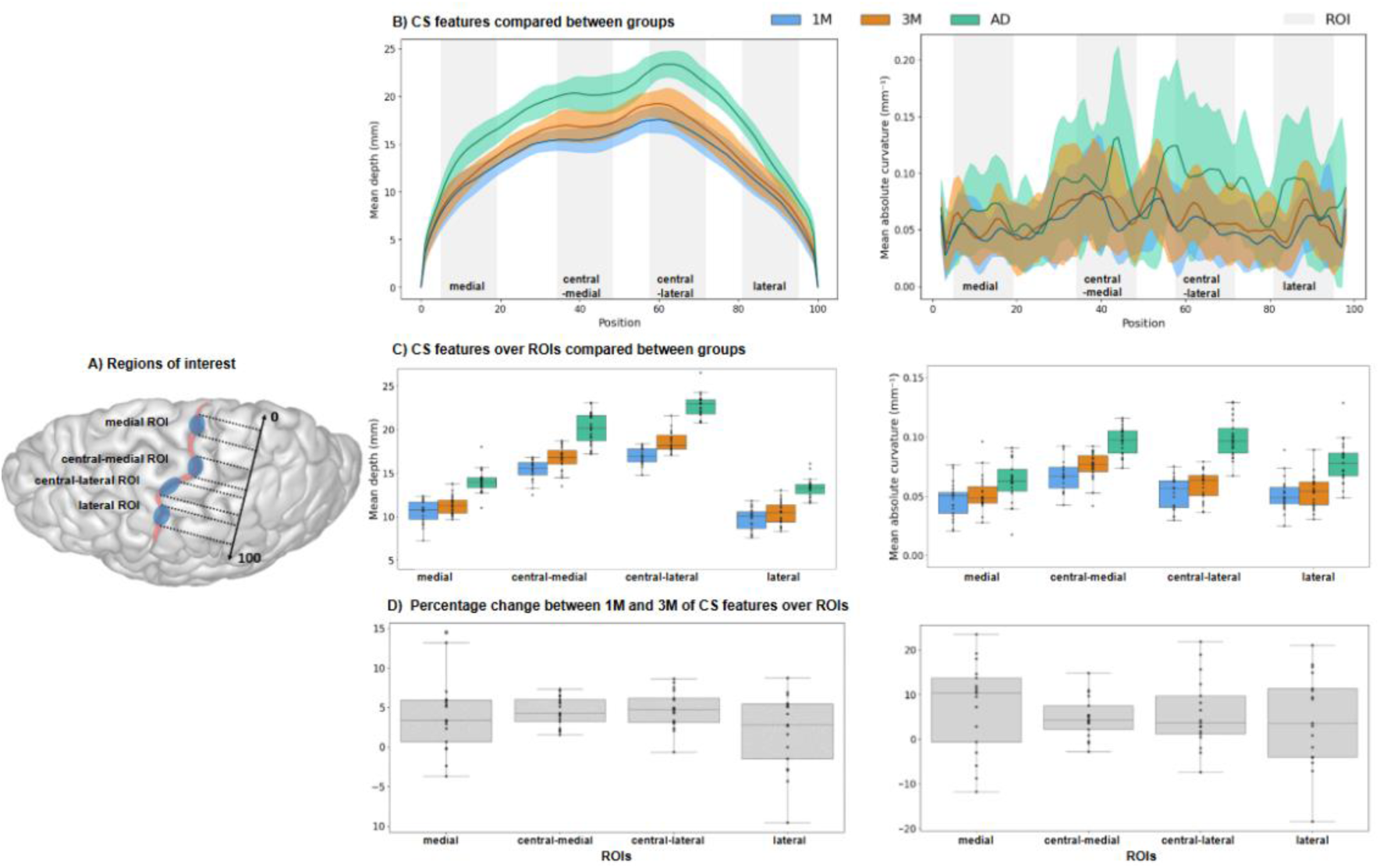
Localization of CS ROIs and feature variability across age groups and ROIs. A) Schema illustrating the ROIs localization along the CS. B) CS parametrization for depth (left column) and absolute curvature (right column), averaged over left and right sulci over all subjects of each group (1M in blue, 3M in orange, AD in green: mean and standard deviation in shaded colours). The averaged position of the 4 ROIs (medial, central-medial, central-lateral, lateral) is highlighted in grey. C) Boxplots of CS depth (left column) and absolute curvature (right column) averaged over each ROI, for the three age groups (same colour code as in B). D) Boxplots of CS feature relative changes between 1M and 3M for infants with longitudinal data.

The LMM model performed on CS depth of the included subjects (1M: N=21, 3M: N=23, AD: N=22) highlighted significant main effects for the factors Group (F(2, 131.50) = 161.66, p < 2.2.10^-16^) and ROI (F(3, 203.75) = 797.70, p < 2.2.10^-16^), as well as a significant interaction between both (F(6, 203.75) = 5.05, p = 3.36.10^-5^). EMMs post-hoc tests (Table 2) first confirmed that depth globally increased with age: CS was significantly deeper at 3M than at 1M in infants, and even deeper in adults than in 3M infants (Figure 4C left column). Second, the four ROIs significantly differed in depth: the central-lateral ROI appeared to be the deepest, followed by the central-medial ROI, the medial ROI, and the lateral ROI appeared to be the shallowest. Third, the ROIs seemed to differently increase in depth with age, as a significant increase between 1M and 3M was only observed for the central-medial and central-lateral ROIs, while a significant increase was observed in all ROIs between 3M and adulthood. Based on the computation of the depth relative difference comparing mean values between 3M and AD groups, we qualitatively observed larger increase in the lateral ROI (12.57%) than in the other ROIs (central-lateral ROI: 10.07%, medial ROI: 10.65%, central-medial ROI: 9.38%).

**Table 2:** Summary of post-hoc statistics in the comparison of depth between Groups and ROIs.

| Contrast |  | Estimate | SE | t.ratio | p.value adjusted |
| --- | --- | --- | --- | --- | --- |
| Effect of Age Group |  |  |  |  |  |
| 1M - 3M |  | -1.08 | 0.19 | -5.76 | < 10-4 |
| 3M - AD |  | -3.28 | 0.24 | -13.71 | < 10-4 |
| Effect of ROI |  |  |  |  |  |
| medial ROI - central medial ROI |  | -5.42 | 0.20 | -26.62 | < 10-4 |
| medial ROI - central lateral ROI |  | -7.44 | 0.20 | -36.49 | < 10-4 |
| medial ROI - lateral ROI |  | 0.86 | 0.20 | 4.21 | 0.0002 |
| central medial ROI - central lateral ROI |  | -2.01 | 0.20 | -9.86 | < 10-4 |
| central medial ROI - lateral ROI |  | 6.28 | 0.20 | 30.83 | < 10-4 |
| central lateral ROI - lateral ROI |  | 8.30 | 0.20 | 40.7 | < 10-4 |
| Effect of Age Group * ROI |  |  |  |  |  |
| Contrast |  | Estimate | SE | t.ratio | p.value adjusted |
| 1M-3M | medial ROI | -0.72 | 0.36 | -2.06 | 0.052 |
|  | central medial ROI | -1.36 | 0.36 | -3.78 | 0.0002 |
|  | central lateral ROI | -1.64 | 0.36 | -4.58 | < 10-4 |
|  | lateral ROI | -0.68 | 0.36 | -1.68 | 0.09 |
| 3M-AD | medial ROI | -2.66 | 0.39 | -6.90 | < 10-4 |
|  | central medial ROI | -3.42 | 0.39 | -8.87 | < 10-4 |
|  | central lateral ROI | -4.13 | 0.39 | -10.7 | < 10-4 |
|  | lateral ROI | -2.93 | 0.39 | -7.61 | < 10-4 |

The LMM model performed on CS absolute curvature of the included subjects (1M: N=21, 3M: N=23, AD: N=21) also revealed significant main effects of Group (F(2,118.81) = 45.33, p = 3.47. 10^-3^) and ROI (F(3,200.95) = 51.59, p = 2.40. 10^-7^), as well as a significant interaction (F(6,200.95) = 6.69, p = 4.24. 10^-7^). EMMs post-hoc tests (Table 3) further revealed the directionality of these effects. As for depth, CS curvature first showed an increase with age: the CS appeared to be significantly curvier at 3M than at 1M, and even curvier in adults (Figure 4C right column). Second, the four ROIs significantly differed in curvature: the central-medial ROI (HK-related) appeared to be the curviest, followed by the central-lateral ROI, the lateral ROI, and the medial ROI was the least curved. Finally, the ROIs displayed distinct patterns of curvature increase throughout development. Between 3M and adulthood, all ROIs exhibited significant changes, except for the medial ROI, which only displayed a tendency towards an increase. No significant changes were observed between 1M and 3M for any of the ROIs, but a trend towards increased curvature was noticed in the central-lateral ROI. Based on the computation of the curvature relative difference comparing mean values between 3M and AD groups, we qualitatively observed larger increase in the central-lateral ROI (23.95%) than in the other ROIs (lateral ROI: 17.91%, central-medial ROI: 12.50%, medial ROI: 8.50%).

**Table 3:**
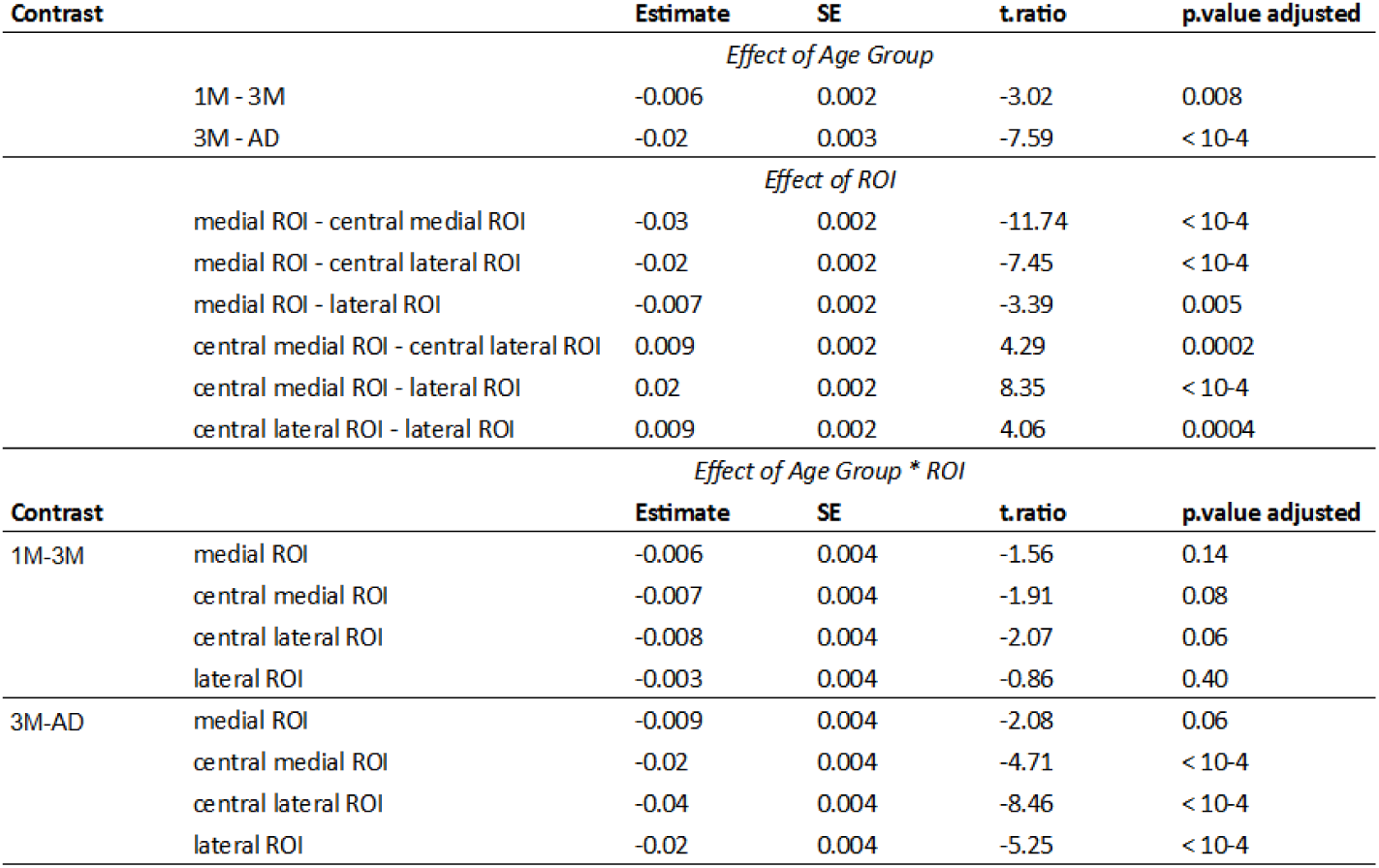
Summary of post-hoc statistics in the comparison of absolute curvature between Groups and ROIs.

| Contrast |  | Estimate | SE | t.ratio | p.value adjusted |
| --- | --- | --- | --- | --- | --- |
| Effect of Age Group |  |  |  |  |  |
|  | 1M - 3M | -0.006 | 0.002 | -3.02 | 0.008 |
|  | 3M - AD | -0.02 | 0.003 | -7.59 | < 10-4 |
| Effect of ROI |  |  |  |  |  |
|  | medial ROI - central medial ROI | -0.03 | 0.002 | -11.74 | < 10-4 |
|  | medial ROI - central lateral ROI | -0.02 | 0.002 | -7.45 | < 10-4 |
|  | medial ROI - lateral ROI | -0.007 | 0.002 | -3.39 | 0.005 |
|  | central medial ROI - central lateral ROI | 0.009 | 0.002 | 4.29 | 0.0002 |
|  | central medial ROI - lateral ROI | 0.02 | 0.002 | 8.35 | < 10-4 |
|  | central lateral ROI - lateral ROI | 0.009 | 0.002 | 4.06 | 0.0004 |
| Effect of Age Group * ROI |  |  |  |  |  |
| Contrast |  | Estimate | SE | t.ratio | p.value adjusted |
| 1M-3M | medial ROI | -0.006 | 0.004 | -1.56 | 0.14 |
|  | central medial ROI | -0.007 | 0.004 | -1.91 | 0.08 |
|  | central lateral ROI | -0.008 | 0.004 | -2.07 | 0.06 |
|  | lateral ROI | -0.003 | 0.004 | -0.86 | 0.40 |
| 3M-AD | medial ROI | -0.009 | 0.004 | -2.08 | 0.06 |
|  | central medial ROI | -0.02 | 0.004 | -4.71 | < 10-4 |
|  | central lateral ROI | -0.04 | 0.004 | -8.46 | < 10-4 |
|  | lateral ROI | -0.02 | 0.004 | -5.25 | < 10-4 |

The analyses of infant’s longitudinal data allowed us to precise these findings by computing the relative rate of change between 1M and 3M at the individual level (Figure 4D). Although longitudinal MRI data were acquired for 22 infants, this analysis focused on 17 of them, after excluding 2 infants with discontinuous CS, 1 outlier in brain volume, and 2 outliers per CS feature. One-sample Student’s t-tests versus zero performed on each ROI over this longitudinal infant group revealed a significant increase of both depth and absolute curvature for all ROIs except for the lateral ROI (Figure 4D and Table 4), suggesting morphological changes of the CS in these three regions between 1M and 3M. However, an ANOVA performed on these relative changes observed across ROIs revealed no significant effect of ROI neither for depth (F(3,64)=1.93, p=0.134, Bayes factor BF10 = 0.56) nor for curvature (F(3,64)=0.34, p=0.799, BF10 = 0.11) suggesting that the evolution of CS features does not differ between regions.

**Table 4.** Summary of Student’s t-tests with the Bayes factor (BF10) statistics examining the relative change between 1 and 3 months of age for each ROI in terms of depth (A) and absolute curvature (B).

| Group | t | relative change (%) | p.value | p.value adjusted | BF10 |
| --- | --- | --- | --- | --- | --- |
| A) Depth |  |  |  |  |  |
| medial ROI | 3.48 | 4.05 | 0.003 | 0.004 | 14.48 |
| central medial ROI | 9.93 | 4.53 | < 10<sup>-4</sup> | < 10<sup>-4</sup> | > 10<sup>+4</sup> |
| central lateral ROI | 8.06 | 4.73 | < 10<sup>-4</sup> | < 10<sup>-4</sup> | 33130 |
| lateral ROI | 1.69 | 1.98 | 0.11 | 0.11 | 0.80 |
| B) Absolute curvature |  |  |  |  |  |
| medial ROI | 2.94 | 7.27 | 0.009 | 0.01 | 5.51 |
| central medial ROI | 4.36 | 4.95 | 0.0004 | 0.002 | 70.57 |
| central lateral ROI | 3.03 | 5.79 | 0.008 | 0.01 | 6.45 |
| lateral ROI | 1.77 | 4.51 | 0.10 | 0.10 | 0.90 |

#### Link to sensorimotor development

For all 29 infants, GM, FM and global motor scaled scores measured with BSID-III evaluations at 3 months of age were in the typical range (Figure 5a). The global score was further analysed according to the evolution of brain volume and CS features in infants with longitudinal measures for both CS depth and absolute curvature (N=15). The first linear model with changes in brain volume showed no significant effect (F(1, 13) = 0.30, r^2^ = 0.02, p = 0.60), nor did the models with the evolution of depth and absolute curvature over the entire CS (F(2, 12) = 2.136, r^2^ = 0.26, p = 0.14; depth: t =-1.54, p = 0.31; curvature: t = 1.14, p = 0.31) and over the central-lateral ROI (F(2,12) = 0.08, r^2^ = 0.01, p = 0.92; depth: t =-0.40, p = 0.85; curvature: t =-0.15, p = 0.97). Although not significant, the model considering the evolution of CS features over the (HK-related) central-medial ROI suggested a trend for a positive relationship between global motor BSID-III scores and the CS depth relative changes between 1M and 3M (F(2,12) = 3.36, r^2^ = 0.36, p = 0.07; depth: t = 2.45, p = 0.12; curvature: t = 0.32, p = 0.72) (Figure 5b). Overall, these results suggest that sensorimotor skills at 3M of age are only weakly related to the early evolution in brain volume and CS morphological features between 1M and 3M of age.

**Figure 5.**
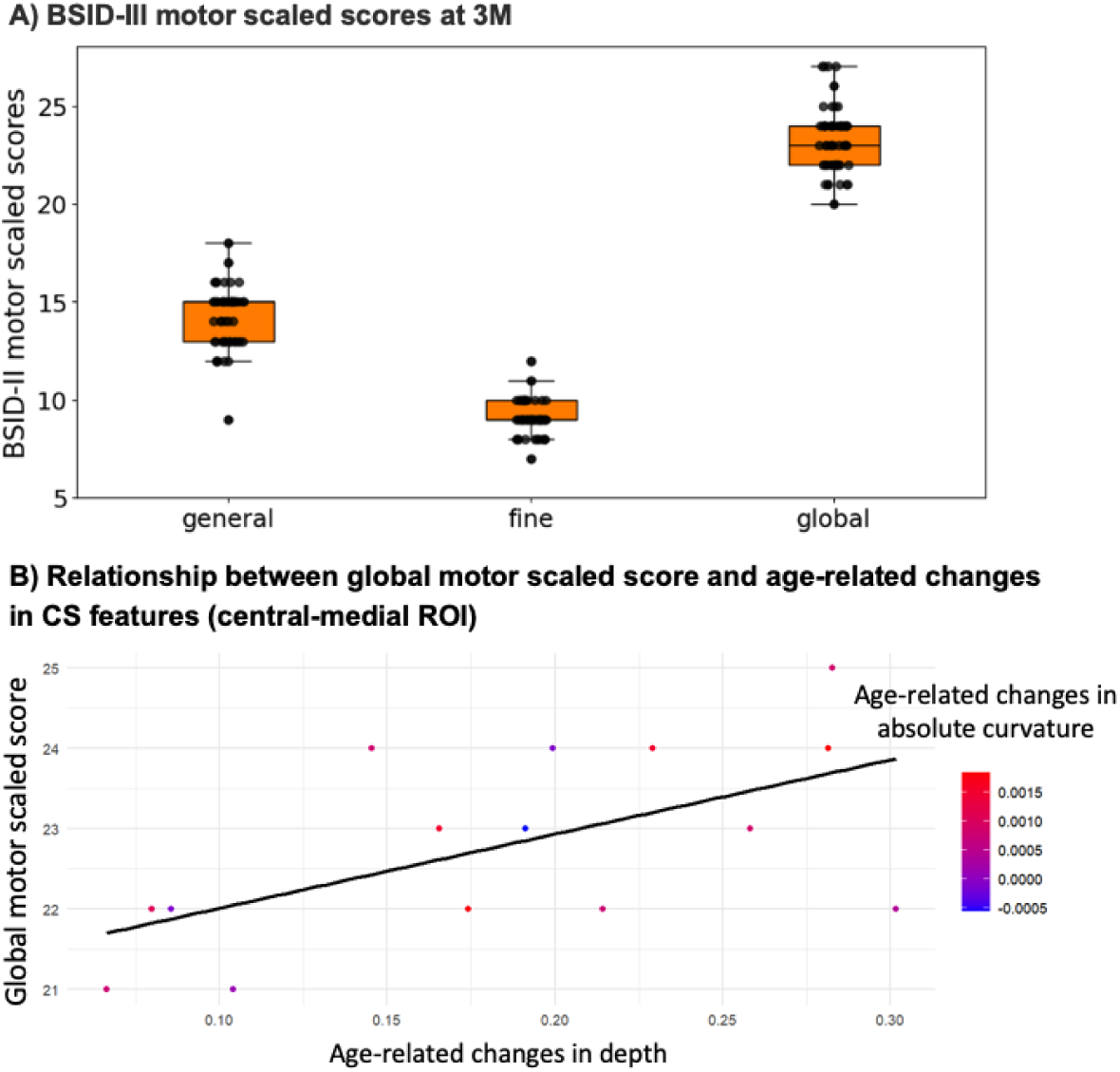
Motor scores at the age of 3 months and relationship with CS morphological changes between 1 and 3 months. A) BSID-III motor scaled scores on the 29 infants at 3 months of age: gross motor (GM), fine motor (FM) and global motor scaled scores. B) Illustration of the linear model testing the relationship between global motor scaled score at 3M and age-related changes in morphological features of the CS central-medial (HK-related) ROI between 1M and 3M (depth represented in abscissa, absolute curvature coded in color).

## Discussion

This study aimed to assess the differential evolution of morphological features along the CS in a unique cross-sectional and longitudinal dataset of typical infants compared to adults with advanced MRI methodologies. We provided evidence for a gradual increase in CS depth and curvature between 1M, 3M and adulthood coherent with the expected increase in brain folding, as well as for morphological differences along the CS across age groups. Interestingly, our cross-sectional results further showed that the increase in CS depth and curvature between 1M and 3M is more pronounced for the central regions of the CS, although longitudinal sub-group analyses failed to detect differences across CS regions. Finally, when evaluating the link to sensorimotor skills, only a trend for a positive relationship between global motor scaled scores at 3M and the depth evolution of the CS central-medial (HK-related) region between 1M and 3M of age was found, suggesting a weak association. Overall, this study provides one of the first quantitative assessment of the evolution and behavioural relevance of CS morphological features in the first months after birth.

### Morphological features of the CS

#### Variations in depth and curvature along the CS

We first highlighted that the non-uniform morphological characteristics along the CS observed in adults are already present in typically developing 1M and 3M-old infants. This arises from a gradient of varying depth and curvature, with the central regions of the CS being the deepest and most curved across all age groups. Regarding CS depth, we observed that the central-lateral ROI was the deepest, consistently with previous studies performed in both adults [30] and 12 and 60M children [10]. The most immediate explanation for a deeper sulcus in central regions arises from the intrinsic construction of the CS sulcal object and the fact that the most medial and lateral regions are by definition close to the anatomical “anchors” that the extremities of the sulcus represent. Although we tried to account for this “anchor” bias by excluding five points on each extremity, this anatomical constrain may limit the observation of depth changes in these regions. It has also been suggested that the depth difference between the central-lateral and central-medial ROIs might rely on the presence of the middle fronto-parietal « pli de passage » (PPFM) [29,30], a privileged place for white matter connections between frontal and parietal lobes, in particular between motor and sensory regions. Showing a higher increase in depth in the central-lateral ROI, our results in 1-3M infants and previous findings in 12-60M-olds [10] seem to be consistent with the development of PPFM and underlying connections in more medial central regions. Besides, we observed that the central-medial ROI showed the most pronounced curvature, likely arising from the omega-shaped hand-knob [32] it includes. While our study was the first to detail CS curvature, we might assume that this higher curvature relies on a more intense cortical development and neuronal proliferation in this region that is further constrained by anatomical anchors such as the PPFM. Overall, we suggest that the consistent variations of CS depth and curvature found across age groups may arise from the combination of anatomical and connectivity constraints in the sulcus definition and development.

#### Inter-hemispheric asymmetry in CS depth and curvature

Another common feature that we observed across age groups is the absence of significant inter-hemispheric asymmetries in CS morphological features. This result might be surprising considering that asymmetries in CS shape were previously reported in adults [6] and in preterm infants imaged at 30 to 40 weeks PMA [7], with a more frequent‘double-knob’ configuration in the right hemisphere compared to the classical’single-knob’ configuration. Similarly, the right CS was found to be deeper than the left in children of 12M of age, particularly in the lateral regions, though this asymmetry disappeared before 24M of age [10]. This discrepancy with our results may arise from the fact that our analysis of asymmetry was performed over the whole CS, which would be consistent with previous findings [34] showing no asymmetry in global CS surface in the same preterm cohort as studied in [7]. Importantly, the adjusted position of our individual ROIs based on the HK location on each hemisphere may prevent us from detecting these asymmetries. In line with this, trends for significant differences in HK location across hemispheres were observed in the 3M and adult groups (see Supplementary Information SI.1 and Supplementary Table 1.1), and no significant asymmetries in depth and absolute curvature were observed in infants when considering the four ROIs (see Supplementary Table 1.2). Overall, our results suggest that while early focal asymmetries might be present in the HK region, there is no difference in overall depth and curvature between the left and right CS in either typical infants or adults.

### Developmental changes in CS morphological features

#### Global increase in CS depth and curvature with age

As anticipated, our results also revealed an increase in CS depth and curvature between 1M and 3M and between 3M and adulthood, coherent with the expected increase in cortical folding. While cerebral development is the result of numerous cellular mechanisms that begin during gestation [35], some authors have proposed the existence of a’folding protomap’ responsible for an early sketch of folding patterns (e.g., [11,36,37]; for a review see [2]). According to this theory, areas of the subventricular zones with higher neuronal proliferation would be responsible for higher expansion of the cortex, giving rise to gyri after neuronal migration [38]. Such mechanism is probably coupled with others to generate the complex dynamics of the folding process, such as the differential growth of brain tissues with different viscoelastic properties, the growth of white matter connections, or the anatomical and physical constraints limiting brain size and connection length to ensure efficient communication between regions (for a review see [2]). Our results are in line with these theories since the CS may become progressively deeper and curvier to adapt to the cortical growth of motor and somatosensory regions in pre-and post-central gyri.

Based on qualitative comparisons of average values over groups, we further observed that the relative difference in CS morphological features was less important than the one in brain volume, both mechanism proceeds earlier than other mechanisms, such as white matter growth that may have a greater impact on brain growth (for a review see [39]). And this may be particularly true for the CS that is known to fold early on [1], in parallel to the early microstructural maturation of the primary sensory and motor regions it delineates (e.g., [20]). Nevertheless, finding significant differences in CS depth and curvature in just two months of development between 1M and 3M suggests that the first post-natal months still represent a critical phase in the folding of early maturing regions, although a deceleration in whole-brain folding was previously observed over this age range compared to the earlier pre-term period [12].

#### Non-uniform changes in depth and curvature along the CS with age

Besides, we observed that the evolution of morphological features with age varied across CS regions. In particular, between 1M and 3M, significant increases in depth were only observed for the central-medial and central-lateral ROIs, as well as only a trend of curvature increase in the central-lateral ROI. This suggests more intense changes in central than medial and lateral ROIs. Several speculations have been made regarding the different stages of CS formation during gestation (e.g., [40]). It would initially be divided into two pieces, the so-called sulcal ‘roots’ or ‘pits’ separated by a cortical bridge (e.g., [41]). As development progresses, the two parts would expand and eventually merge, creating a fold where the cortical bridge initially separated them. Once this fusion is complete, development appears to continue primarily at the centre of the emerging sulcus, where the fold formed during fusion further widens and deepens, gradually making the cortical bridge disappear from the surface. Our cross-sectional results of increased depth in the two CS central regions, but not in medial and lateral regions, support this model, highlighting significant early changes occurring specifically in central regions. These are also in agreement with the most intense changes reported in CS central regions in 12 to 60M old children [10]. Nevertheless, our analysis of longitudinal data did not confirm the difference in depth increase across ROIs, perhaps because of the limited sample size and/or to the higher variability in depth measures in the medial and lateral ROIs.

Besides, it is important to note that within our initial group of 33 infants, 3 had a discontinuous CS. According to a recent theory [11], the formation of a discontinuous CS might result partly from a premature development of the cortical bridge (or PPFM) connecting the pre-and post-central gyri, in comparison to the development of the two independent pieces of the sulcus. However, given the rarity of these cases observed in adults – 6 cases observed out of 2174 hemispheres in a previous study [42] – and the frequency of these cases observed within our infant sample, we might assume that the CS discontinuities we detected are not necessarily permanent but that CS folding, particularly at its centre with PPFM development, is still ongoing during the first months after birth. Scanning those 3 infants at a later age would allow us to test this hypothesis.

Aside from anatomical substrates, an intense development of CS central regions during infancy may also be related to differential functional evolution of sensorimotor regions related to different body parts. The central-medial and central-lateral ROIs are thought to correspond to the upper limbs and face which may develop more intensely at these ages since they are most densely innervated [5] and are intensely co-stimulating each other from in-utero ([43], for a review see [9]). In line with this, a recent functional MRI study on preterm infants [44] provided evidence for the existence of a somatotopic organization at 32-weeks PMA, already displaying a higher magnification for the hand region than for the ankle or the mouth. Nevertheless, it would be informative to confirm our hypotheses by linking CS morphological features with the developing somatotopic organization in typical infants, based on a functional mapping of different body parts.

#### Methodological considerations

A first consideration arose from the fact that age-related increases and differences across regions were more systematically detected with CS depth than with curvature. While curvature could be expected to develop in parallel to depth, it may be a marker more difficult to quantify or less sensitive to the cortical expansion process over this developmental period due to more stable mechanical constraints, limiting its progression compared to depth. In addition, averaging over rather large ROIs may have prevented us from detecting small and more local changes in curvature with age, though a point-to-point analysis along the CS did not reveal stronger effects (see Supplementary Information SI.2 and Supplementary Figure 2.1). Finally, the higher heterogeneity and inter-individual variability (see Supplementary Information SI.3) of this feature may make it more difficult to detect group differences.

Another consideration comes from the fact that cross-sectional and longitudinal analyses were not fully consistent to detect differences across CS regions. Here again, the high inter-individual variability present in developmental populations (each infant following a specific neurodevelopmental trajectory) may have contributed to differences observed between cross-sectional and longitudinal analyses. We tried to limit this variability by defining the ROIs at the individual level, rescaling them to each infant and age, as well as positioning them relatively to the position of the hand-knob and CS extremities. Nevertheless, we cannot confidently assert that the different ROIs evolve in a homothetic manner between 1M and 3M, which may limit their anatomical correspondence relative to the somatotopic organization and thus impede their comparison between infants and across ages.

Another methodological limitation of our study arose from the intrinsic variability of the pipeline used for the sulcal object identification, particularly at CS extremities where the lower depth led to a noisier and more variable identification of CS object. While this variability appeared to have a relatively minor impact on our results (see Supplementary information SI.3), it is important to emphasize its potential implications for comparisons between subjects and age groups.

### Relating the development of CS morphological features and sensorimotor skills

The last objective of our study was to relate the developmental changes in CS morphological features with the acquisition of sensorimotor skills in typical infants. Although all tested models revealed non-significant results, a trend of relationship was observed for the age-related increase in the depth of the (HK-related) central-medial ROI. This trend may be related to the increased hand usage that was found between 1 and 3M [45]. At the beginning of life, infants have limited means to explore the world, which lead them to adopt mostly self-touch behaviours (for a review, see [9]). They begin touching their face during the foetal stage [43] and continue after birth, initially making contact with their head, chest, and hands, before gradually moving towards their hips and feet [13]. Previous studies seem to show an increasing usage of the hands between 1 and 3 months, progressing from self-touch behaviour to the emergence of reaching and object manipulation around 3 to 5 months [45]. While impossible to detect with Bayley scales at such early ages, this increasing hand usage may relate to the CS depth maturation in the (HK-related) central-medial ROI.

The absence of observed relationship between CS anatomical and behavioural development could be explained by various limitations. First, the sample size was limited to 15 infants as analyses required longitudinal measures for both CS depth and curvature, requiring the exclusion of all outlier infants. Second, though it is more readily available in young infants (with a short MRI acquisition achievable during natural sleep), the CS morphology might be a too indirect marker of the development of sensorimotor regions. It would be interesting to test the shape analysis approach [6], which had shown certain relationships between the CS shape of preterm infants and motor development at 5 years [7]. Measuring the microstructural maturation of the pre-and post-central cortices [20] would be an interesting way of obtaining more specific markers.

Another limitation comes from the battery of tests we used to assess global motor skills in infants at 3M (Bayley scale, BSID-III; [21]). Although this scale is frequently used in young children to detect neurodevelopmental disorders (e.g., [46,47]), it may lack sensitivity in young infants, in particular to assess fine motor skills. Indeed, assessment items at 3M focus more on postural control (e.g., holding the head in different positions) and on fine motor skills that involve gaze evaluations than on hand and mouth usage (e.g., grasping a ring or hand-to-mouth behaviour). This neurodevelopmental evaluation further depends on the infant state of alertness, readiness and fatigue, but we were particularly careful to carry out the experiments under similar conditions. Importantly, previous studies relating early CS development and motor skills in preterm infants actually considered outcomes at later ages: at either 24 or 30 month of age (corrected for gestational age at birth) with BSID-III [34], or at 5 years of age with the Movement Assessment Battery for Children, 2^nd^ edition (MABC-2) [7]. Following our typical infants at later ages may thus be important to get a more relevant picture of their neurodevelopmental trajectory. Nevertheless, the trend in the relationship observed with the CS region associated with the hand may make us optimistic to continue exploring this avenue of research.

## Conclusion

Relying on a unique cross-sectional, longitudinal and multimodal design to study typical infants, we could capture the progressive evolution of CS morphological features across regions supposed to relate to somatotopic organization. The most intense and robust changes were observed in the central regions, including the hand-knob which was the only region to show some trend of relationship with the acquisition of sensorimotor skills. Such framework might help identify early alterations in neurodevelopmental trajectories and design early interventions to improve the outcome of infants at risk of developing sensorimotor disorders.

## Abbreviations

AD: adult
AI: asymmetry index
BSID-III: Bayley Scales of Infant and Toddler Development, 3rd edition
CS: central sulcus
EMMs: estimated marginal means
FDR: False Discovery Rate
GM / FM: gross / fine motor
HCP: Human Connectome Project
HK: hand knob
L / R: left / right
LMM: linear mixed-effects model
M1: primary motor cortex
MRI: magnetic resonance imaging
PMA: post-menstrual age
PPFM: middle fronto-parietal « pli de passage »
ROI: region of interest
S1: primary somatosensory cortex
T1w / T2w: T1 / T2-weighted
1M / 3M: 1 month / 3 month**s**

## Acknowledgement

The authors thank Marianne Barbu-Roth’s team (Marie-Victorine Dumuids-Vernet, Léa Guéret, Viviane Huet and Joëlle Provasi) for their advices on the BSID-III scale, David Germanaud for his advice on infants inclusion criteria, Denis Rivière, Olivier Coulon and Héloïse de Vareilles for discussion on the central sulcus analyses and parameterization, Laurie Devisscher for organizing HCP data, the UNIACT team for the experiment organization at NeuroSpin (including the nurses and MRI technologists, and Bernadette Martins for ethical questions), and for their valuable help in recruiting infants: Christine Doublé, Marie Palu and healthcare professionals, including the team of midwives and pediatricians (Jean Gaschignard and Sophie Gobet) at the Orsay maternity hospital. And, last but not least, the authors would like to sincerely thank all the infants and parents who participated in this study, as well as adult participants.

## Statement of Ethics

### Study approval statement

This study was performed with the agreement of the national ethical committee “Comité de Protection The study design and procedures was reviewed and approved by the ethics committee (Comité de Protection des Personnes, CPP Ile de France 3), under the protocol DEVine (CEA 100 054; ID RCB 2020-A00106-33) for infants, and the protocol Methodo3T (CEA 100 052; ID RCB 2019-A02482-55) for adults.

### Consent to participate statement

Parents were informed about the study and gave written consent to participate with their babies. Adult participants also provided written informed consent to the study.

## Author Contributions

Conceptualization: JD, DM, MBR, JFM

Data curation: AD, DM, AG

Formal analysis: AD, DM

Funding acquisition: JD, MBR, JFM

Investigation: DM, AG, LHP

Methodology: AD, YL, DM, JD

Project administration: JD, DM, MBR

Resources: YL, JFM

Supervision: DM, JD

Validation: DM, JD

Visualization: AD, DM

Writing—original draft: AD, DM, JD

Writing—review & editing: all.

## Data Availability Statement

The data that support the findings of this study are not publicly available due to their containing information that could compromise the privacy of research participants but are available from JD upon reasonable request.

