## Supplementary Information for "On the typical development of the central sulcus in infancy: a longitudinal evaluation of its morphology and link to behaviour"

### SI.1: Evaluation of inter-hemispheric asymmetries

#### *Method*

We first evaluated whether the position of the hand knob (HK) along the CS differed between the left and right hemispheres. Wilcoxon signed-rank tests were conducted on asymmetry indices (AI) of HK coordinates for each age group.

Secondly, to complement our findings across the entire central sulcus (CS) (main manuscript), we evaluated whether our regions of interest (ROIs) showed inter-hemispheric asymmetries in CS morphological features. For each group and each ROI, we computed individual AI of depth and absolute curvature. The presence or absence of asymmetry was evaluated for each age group independently using Wilcoxon test on AI.

Corrections for multiple comparisons were performed using the false discovery rate (FDR) approach.

#### *Results*

Regarding HK position, the Wilcoxon signed-rank tests performed on each age group showed no significant differences between the left and right hemispheres for 1M and 3M infant groups, but an almost significant asymmetry was observed for the adult group, confirmed by Bayes factor (BF10 > 1) (**Supplementary Table 1.1**). These results suggest a tendency towards an inter-hemispheric asymmetry in the HK position, becoming more pronounced with age.

***Supplementary Table 1.1:*** *Results of the Wilcoxon signed-rank tests on asymmetry indices for the position of the hand-knob for the 1M, 3M, and AD groups. Legend: N, number of subjects per group; BF10, Bayes factor.*

| Group | N | W | p.value | p.value adjusted | BF10 |
|---|---|---|---|---|---|
| 1M | 23 | 72 | 0.22 | 0.22 | 0.60 |
| 3M | 25 | 87 | 0.12 | 0.18 | 0.91 |
| AD | 23 | 48 | 0.02 | 0.06 | 3.42 |

Regarding asymmetries in depth and absolute curvature over CS ROIs, the Wilcoxon tests showed no significant results for any age group, except for the lateral ROI in adults, where a leftward asymmetry in the CS was observed (**Supplementary Table 1.2**).

***Supplementary Table 1.2:*** *Results of the Wilcoxon tests on asymmetry indices for depth (A) and absolute curvature (B) averaged over ROIs, for the 1M, 3M, and AD groups.*

| Group by ROI | | N | median | W | p.value.adjusted | BF10 |
|---|---|---|---|---|---|---|
| **A) Depth** | | | | | | |
| 1M | medial ROI | 21 | 0.0006 | 114 | 0.97 | 0.23 |
| | central medial ROI | 21 | 0.018 | 106 | 0.97 | 0.23 |
| | central lateral ROI | 21 | -0.005 | 80 | 0.92 | 0.46 |
| | lateral ROI | 21 | 0.004 | 99 | 0.97 | 0.25 |
| 3M | medial ROI | 23 | 0.009 | 102 | 0.44 | 0.34 |
| | central medial ROI | 23 | 0.01 | 115 | 0.73 | 0.25 |
| | central lateral ROI | 23 | -0.02 | 95 | 0.32 | 0.34 |
| | lateral ROI | 23 | 0.02 | 105 | 0.50 | 0.31 |
| AD | medial ROI | 22 | 0.02 | 105 | 0.95 | 0.25 |
| | central medial ROI | 22 | -0.002 | 124 | 0.95 | 0.23 |
| | central lateral ROI | 22 | 0.004 | 117 | 0.95 | 0.23 |
| | lateral ROI | 22 | 0.03 | 70 | 0.06 | 1.37 |

| **B) Absolute curvature**<br>Group by ROI | | N | median | W | p.value.adjusted | BF10 |
|---|---|---|---|---|---|---|
| 1M | medial ROI | 21 | 0.008 | 105 | 0.95 | 0.26 |
| | central medial ROI | 21 | -0.004 | 101 | 0.95 | 0.31 |
| | central lateral ROI | 21 | -0.02 | 113 | 0.95 | 0.24 |
| | lateral ROI | 21 | -0.02 | 108 | 0.95 | 0.25 |
| 3M | medial ROI | 23 | 0.01 | 102 | 0.22 | 0.37 |
| | central medial ROI | 23 | 0.04 | 121 | 0.14 | 0.24 |
| | central lateral ROI | 23 | 0.05 | 85 | 0.22 | 0.71 |
| | lateral ROI | 23 | -0.05 | 115 | 0.3 | 0.24 |
| AD | medial ROI | 21 | -0.11 | 96 | 0.66 | 0.25 |
| | central medial ROI | 21 | -0.02 | 100 | 0.66 | 0.24 |
| | central lateral ROI | 21 | 0.0004 | 102 | 0.66 | 0.27 |
| | lateral ROI | 21 | 0.10 | 44 | 0.045 | 12.54 |

### SI.2: Analyses of depth and absolute curvature along the CS

#### *Method*

In addition to studying the evolution of depth and curvature in specific ROIs along the sulcus, we also aimed to globally analyze how these morphological metrics evolve at every CS position between infants at 1M and 3M and adults. To achieve this, we performed Mann-Whitney tests to compare depth and absolute curvature measurements (averaged over both hemispheres) across groups (1M vs 3M, 3M vs adults) for each coordinate along the CS (from 5 to 95). Multiple comparisons were corrected using permutation testing, following the approach previously described [1].

#### *Results*

Global analyses of the age-related changes in depth and absolute curvature along the CS are presented in **Supplementary Figure 2.1**. For CS depth, Mann-Whitney tests revealed significant differences between groups in most CS positions: depth was higher in 3M than in 1M infants in positions 15-66 and 74-75 (corrected $p < 0.05$), and in all positions in adults compared with 3M group (corrected $p < 10^{-4}$). These findings are consistent with our main results showing larger depth changes between 1 and 3M of age in central ROIs than in medial or lateral ROIs, and in all ROIs between 3M and adult groups (Table 2).

In contrast, for absolute curvature, the analysis along the CS with Mann-Whitney tests revealed fewer significant group differences than for depth: no difference between 1M and 3M, and only a few positions showed higher curvature in adults that 3M infants (corrected $p < 0.05$ for positions 43-46, 56-59, 65-75, 83-86 corresponding to the central and lateral parts of the sulcus). These results are relatively consistent with our main ROI analysis (Table 3) showing a trend of change between 1M and 3M only for the central-lateral ROI, and changes between 3M and adult groups for central and lateral ROIs in addition to a trend for the medial ROI. This suggests that absolute curvature is a feature showing high variability along the sulcus, and that differences between groups are more difficult to demonstrate point-to-point than by ROIs.

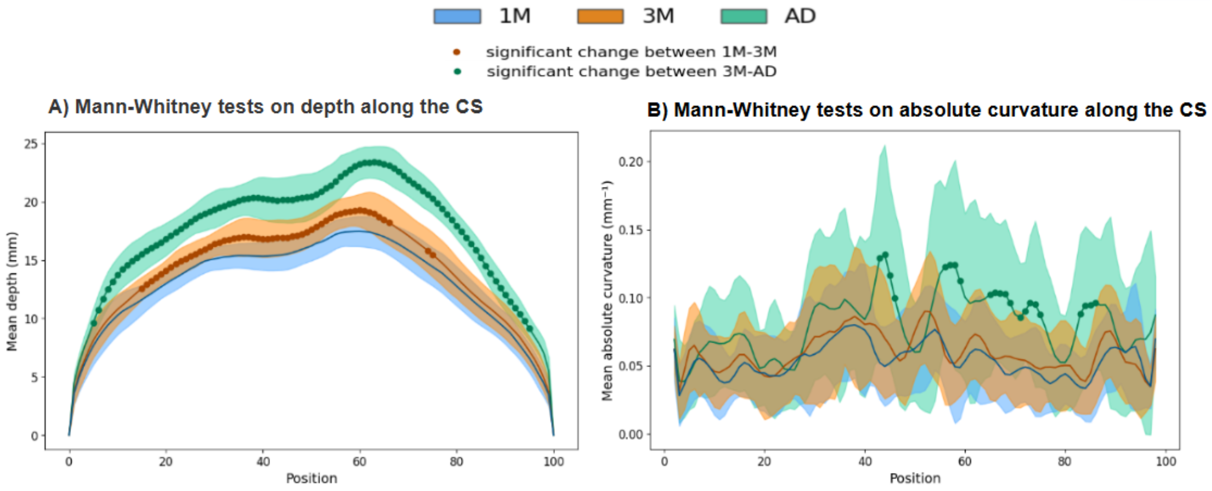


***Supplementary Figure 2.1. Age-related changes in depth and absolute curvature along the CS***
*Profiles of depth (A) and absolute curvature (B) along the CS, averaged over each group (1M, 3M, AD), and showing positions (between 5 and 95) where significant differences between two age groups were observed with Mann-Whitney tests (dots: p<0.05 after permutation testing).*

### SI.3 Evaluation of variability related to methodological approaches

#### *Method*

Additional analyses were conducted to ensure that the software variability did not interfere with the different effects we observed.

#### *Evaluation of intra- vs. inter-subject variability*

First, we considered the variability related to CS identification with Morphologist pipeline and manual labeling. To evaluate intra-subject variability, repeated analyses were performed on three infants with longitudinal data: Subject A (10 repetitions), Subject B (10 repetitions), and Subject C (5 repetitions), at both 1M and 3M and focusing on the left CS. Each analysis was conducted using identical brain segmentations and input parameters, differing only in the independent execution of the pipeline for CS identification. For each repetition, the left CS was identified, and measurements of depth and absolute curvature along it were extracted. For each morphological feature (depth, absolute curvature), we then computed intra-subject cross-correlations along the CS, for all possible pairs of repetitions within each subject (45 pairs for Subjects A and B, 10 pairs for Subject C). For each subject, the mean and standard deviation of the resulting cross-correlation coefficients were computed to characterize intra-subject variability.

Subsequently, we quantified inter-subject variability with a similar approach. For each age (1M, 3M), we selected a given CS identification/analysis from the repeated analyses for each Subject A, B, C, along with a single analysis from 12 additional subjects, all processed using the same pipeline. For depth and absolute curvature respectively, the inter-subject cross-correlations were computed between all possible pairs among these 15 analyses. The mean and standard deviation of the resulting 105 cross-correlation coefficients were computed to characterize inter-subject variability.

Qualitative descriptions of these analyses were done to compare intra- and inter-subject variability, as well as variability in depth and absolute curvature. Because repeated analyses in Subjects A, B, C were performed at both ages, we further evaluated whether intra-subject variability differed between 1M and 3M using Student t-tests on cross-correlations coefficients for depth and absolute curvature respectively.

#### *Evaluation of variability related to image reconstruction: resampling vs. super resolution*

In this study, T2-weighted (T2w) anatomical images were acquired in the axial plane for all infants, and in sagittal plane for a sub-part of them (N=20 across the 1M and 3M groups). To obtain similar isotropic spatial resolution (0.8mm) across all infants for sulcus analysis, we resampled axial T2w images with Fourier interpolation. Here we aimed to evaluate whether this resampling approach may impact CS depth and curvature measurements compared to those obtained when considering images with higher raw resolution. For one infant (Subject A) at both ages (1M, 3M), we reconstructed T2w images with a super-resolution (SR) algorithm (tool Niftymic [2]), based on axial and sagittal T2w volumes and leading to a 0.8mm resolution. As for the processing of resampled axial T2w images, segmentation was further performed using iBEAT2 (version 120, [3]) and manually corrected if necessary using BrainVISA software. CS was identified, and its depth and absolute curvature analyzed. We compared the measurements obtained from SR images with the ones from resampled images: given the inherent intra-individual variability associated with the pipeline, SR-related measurements were compared with the 10 intra-subject measurements obtained with resampled images (10 cross-correlations), for each age and each morphological feature separately. Mean and standard deviation over the 10 resulting cross-correlation coefficients were computed to assess the consistency of the results based on the image reconstruction approach.

Subsequently, Student t-tests on cross-correlation coefficients were performed at each age and for each feature to determine whether the differences observed between SR and resampling approaches were significantly greater than the differences observed between the 10 repeated analyses with resampling approach (45 cross-correlations characterizing intra-individual variability for Subject A). We also evaluated whether the differences observed between SR and resampling approaches differed between age groups. Corrections for multiple comparisons were performed using the FDR approach.

### *Results*

#### *Intra- vs. inter-subject variability*

Repeated analyses for each subject showed relatively high consistency in depth and absolute curvature along the CS at both ages (**Supplementary Figure 3.1.A**). All conducted intra-subject cross-correlations showed relatively high consistency (coefficients between ~0.8 and 1 for intra-individual variability), although some differences were observed across morphological features and across age groups (**Supplementary Table 3.1**, **Supplementary Figure 3.1.B**). Indeed, intra-subject consistency appeared qualitatively higher for depth than for absolute curvature (mean cross-correlation coefficients of individuals ranging from 0.97-0.99 for depth, and 0.87-0.93 for curvature). And the intra-individual consistency was significantly lower at 3M than 1M for depth but not for absolute curvature (results of Student t-tests in **Supplementary Table 3.2**). Besides, for both depth and absolute curvature at both ages, the intra-subject consistency clearly appeared higher (i.e., higher coefficients) than the inter-subject consistency as illustrated in **Supplementary Figure 3.1.B**. This suggests that, despite some intra-subject variability observed in the CS feature measurements, our pipeline demonstrated sufficient reliability not to undermine the effects we aimed to demonstrate in our main analyses across different subjects (main effects and interaction of age group and ROI).

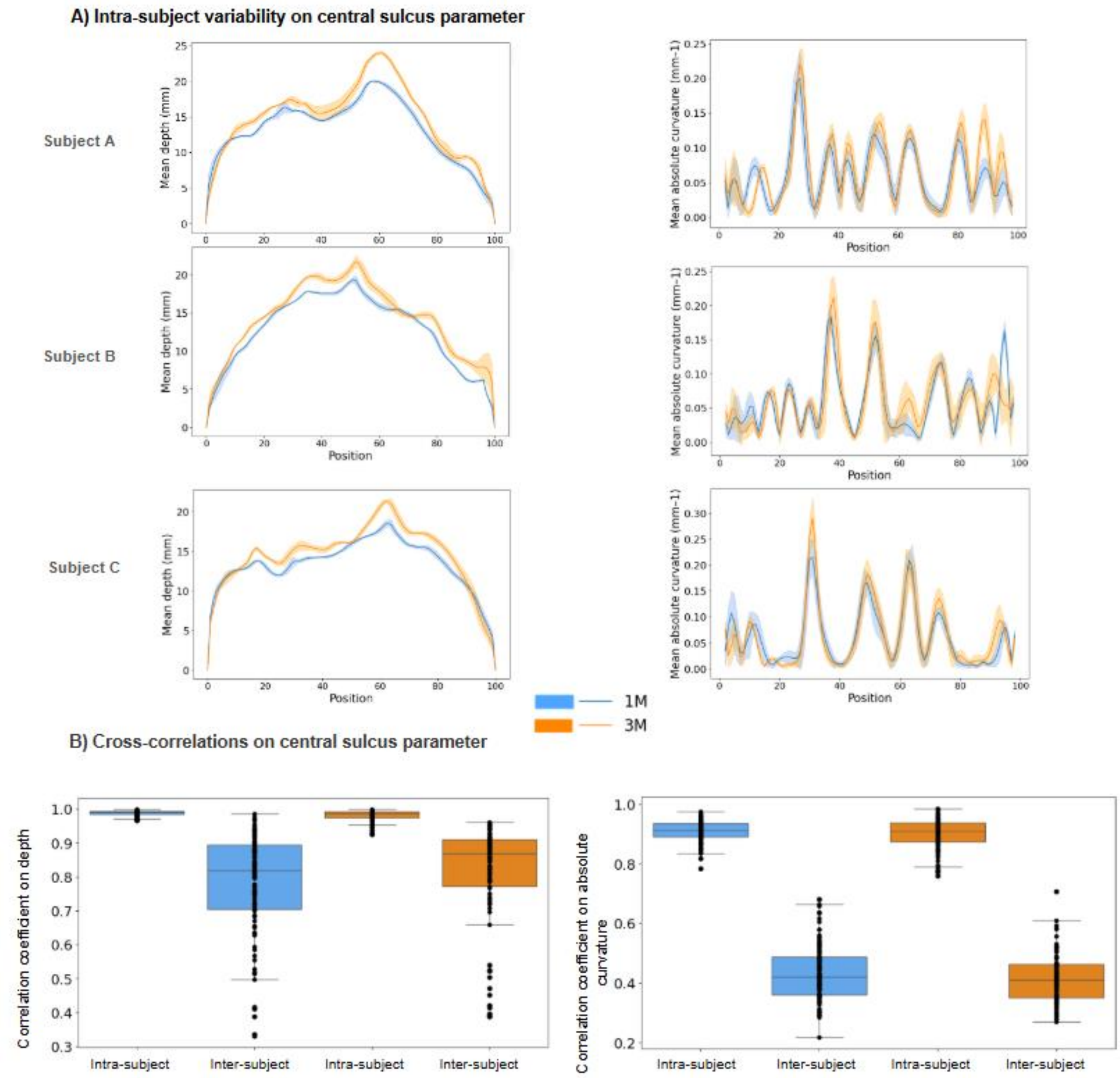


***Supplementary Figure 3.1. Evaluation of intra-subject vs. inter-subject variability.***

*A) Mean and standard deviation of depth and absolute curvature along the CS on repeated analyses performed on Subject A, B and C at 1M and 3M. B) Cross-correlation coefficients for intra-subject (repeated analyses on Subject A, B and C) and inter-subject (analyses on 15 different subjects) at 1M and 3M, on depth and absolute curvature (analyses with resampling approach).*

***Supplementary Table 3.1:*** *Intra-subject cross-correlations for depth (A) and absolute curvature (B) (analyses with resampling approach). Mean and standard deviation were computed over pairs of comparisons (N), for each of the three subjects of interest at 1M and 3M.*

| Group | | N | mean | SD |
|---|---|---|---|---|
| | | *Intra subject cross-correlations* | | |
| 1M | Subject A | 45 | 0.99 | 0.006 |
| | Subject B | 45 | 0.98 | 0.008 |
| | Subject C | 10 | 0.99 | 0.002 |
| 3M | Subject A | 45 | 0.99 | 0.01 |
| | Subject B | 45 | 0.97 | 0.02 |
| | Subject C | 10 | 0.99 | 0.006 |
| **B) Absolute Curvature** | | | | |
| 1M | Subject A | 45 | 0.90 | 0.04 |
| | Subject B | 45 | 0.92 | 0.02 |
| | Subject C | 10 | 0.88 | 0.05 |
| 3M | Subject A | 45 | 0.88 | 0.05 |
| | Subject B | 45 | 0.93 | 0.04 |
| | Subject C | 10 | 0.87 | 0.04 |

***Supplementary Table 3.2****: Results of Student t-tests on intra-subject cross-correlation coefficients between 1M and 3M for depth (A) and absolute curvature (B) (analyses with resampling approach).*

| Contrast | t | p.value.adjusted | BF10 |
|---|---|---|---|
| **A) Depth** | | | |
| 1M-3M | 4.22 | 0.0001 | 321.33 |
| **B) Absolute curvature** | | | |
| 1M-3M | 1.25 | 0.21 | 0.24 |

***Variability related to the image reconstruction approach: super-resolution vs resampling***

Visually, the SR approach provided clearer visualization of the complex folding patterns on T2w images (i.e., less smoothness) than the resampling approach, but nevertheless brain tissues appeared correctly segmented with both approaches (see example of Subject A at 1M in **Supplementary Figure 3.2**).

For this specific subject, the depth and absolute curvature obtained along the CS with the SR approach (dashed lines in **Supplementary Figure 3.3.A**) were relatively in the variability range observed across repetitions with the resampling approach (solid lines and shaded area in **Supplementary Figure 3.3.A**), despite clear differences in some positions, particularly for curvature measurements at the lateral CS extremity. This was confirmed by the high cross-correlation coefficients obtained between the SR and "resampling repetitions" approaches at both ages for CS depth measurements (coefficients > 0.95), revealing a similar range of values than obtained when assessing the intra-subject variability of our pipeline on resampling repetitions (**Supplementary Figure 3.3.B**). For the absolute curvature, the results of such comparisons were more variable: cross-correlations between the SR and resampling repetitions approaches showed systematically lower coefficients than cross-correlations between resampling repetitions (**Supplementary Figure 3.3.B**). T-tests comparing the variability related to the reconstruction approach (cross-correlations between SR and resampling repetitions vs. cross-correlations of resampling repetitions) confirmed significant differences at both ages for the absolute curvature but also for depth at 1M, the variability being lower for resampling repetitions (**Supplementary Table 3.3**). This suggests that the variability related to the reconstruction approach is higher than the variability related to the sulcus pipeline itself.

Looking at potential age-related differences, SR-resampling coefficients appeared relatively higher at 1M (coefficients > 0.8) than at 3M (coefficients > 0.5), something confirmed with Student t-tests for the absolute curvature but not for depth (**Supplementary Table 3.4**). This suggests that our pipeline of CS reconstruction may struggle to properly capture CS curvature more at older ages without having high-resolution images.

Further analyses on additional subjects and with more repetitions would be needed to confirm these observations and decipher whether the SR approach would be more robust to extract CS morphological features.

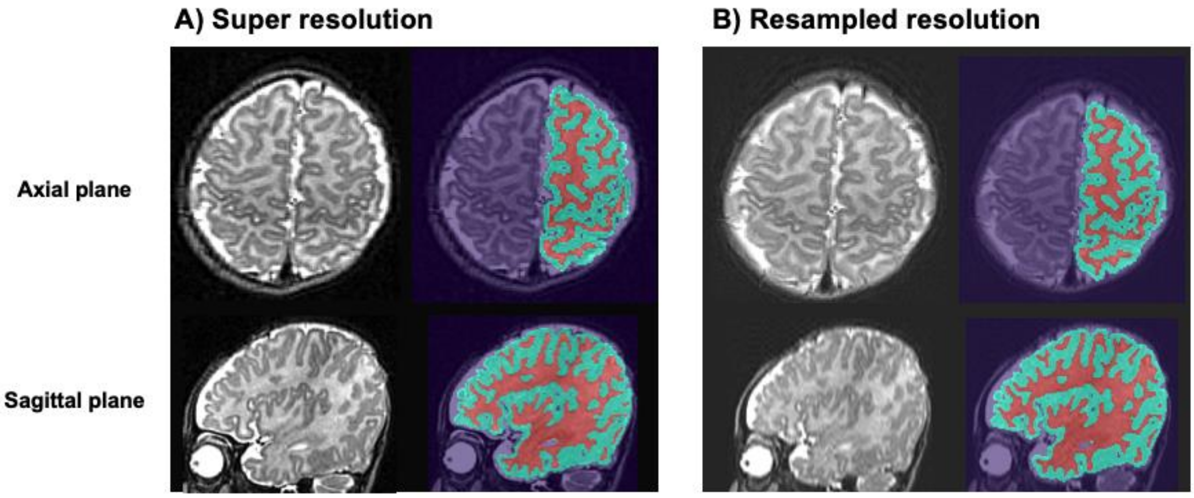


***Supplementary Figure 3.2: Comparison of T2w images and brain tissue segmentations between reconstruction approaches.***
*Example from an individual infant (Subject A at 1M) of the axial and sagittal views of T2w images (left columns) and brain tissue segmentation (right columns: cortical grey matter in green, white matter in red) obtained at the level of the central sulcus, for the approaches of super-resolution (A) and resampling (B).*

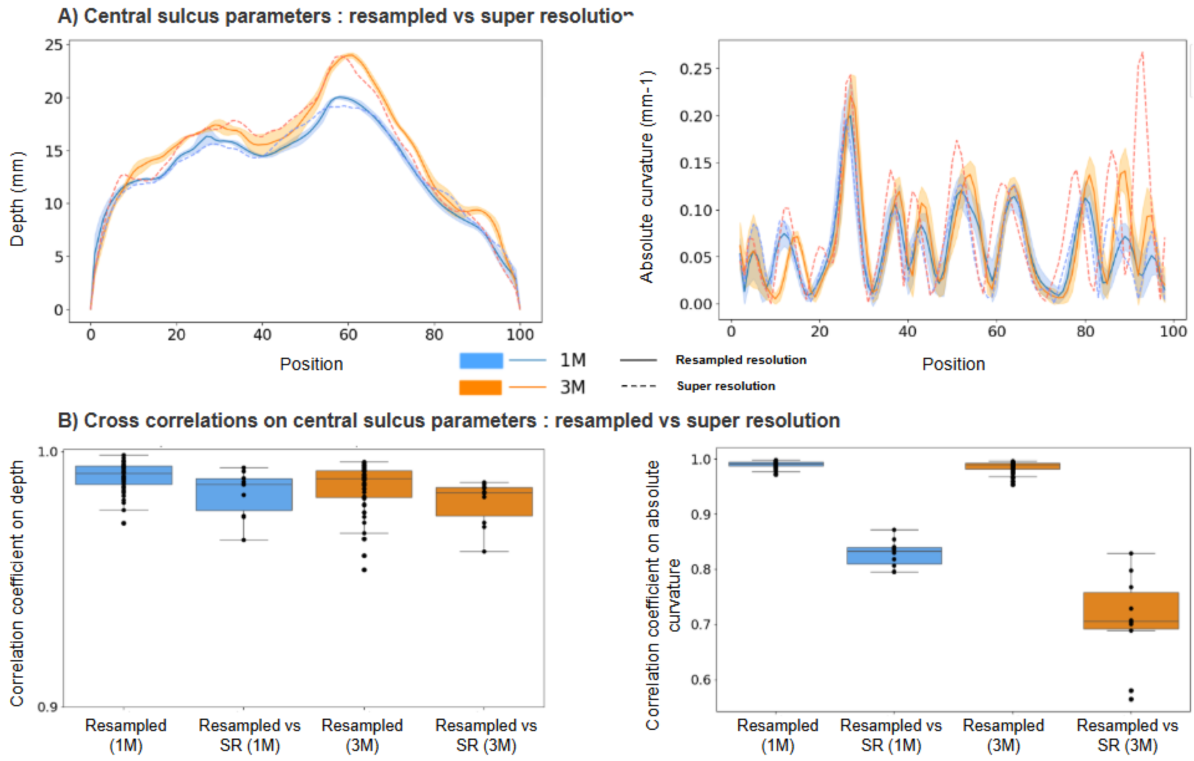


***Supplementary Figure 3.3. Evaluation of reconstruction approaches: super-resolution vs. resampling.*** *A) Depth and absolute curvature along the CS extracted from the SR approach of Subject A at both ages (dashed lines), compared to the mean and standard deviation obtained from the repeated resampled analyses of the same infant (solid lines and shaded area). B) Cross-correlations between the SR and the 10 resampling repetitions approaches, compared to the cross-correlations between the 10 resampling repetitions, for depth (left) and absolute curvature (right) of Subject A at 1M and 3M.*

***Supplementary Table 3.3:*** *Results of Student t-tests comparing the intra-subject variability related to the reconstruction approach (cross-correlation coefficients for SR vs. resampling repetitions) and related to the pipeline (cross-correlation coefficients for resampling repetitions), for depth (A) and absolute curvature (B) of subject A at 1M and 3M.*

| Contrast | t | p.value.adjusted | BF10 |
|---|---|---|---|
| **A) Depth** | | | |
| 1M | -2.94 | 0.006 | 2.00 |
| 3M | -1.7 | 0.09 | 1.14 |
| **B) Absolute curvature** | | | |
| 1M | -40.24 | <10-4 | > 10+4 |
| 3M | -22.19 | <10-4 | >10+4 |

***Supplementary Table 3.4:*** *Results of Student t-tests comparing cross-correlation coefficients for the SR vs. resampling repetitions approaches between 1M and 3M, for depth (A) and absolute curvature (B) of subject A.*

| Contrast | t | p.value.adjusted | BF10 |
|---|---|---|---|
| **A) Depth** | | | |
| 1M-3M | 0.88 | 0.40 | 0.43 |
| **B) Absolute curvature** | | | |
| 1M-3M | 3.82 | 0.008 | 12.78 |
